# The hidden structural variability in avian genomes

**DOI:** 10.1101/2021.12.31.473444

**Authors:** Valentina Peona, Mozes P. K. Blom, Carolina Frankl-Vilches, Borja Milá, Hidayat Ashari, Christophe Thébaud, Brett W. Benz, Les Christidis, Manfred Gahr, Martin Irestedt, Alexander Suh

**Author notes:** Corresponding author: VP; AS.

## Abstract

Structural variants (SVs) are DNA mutations that can have relevant effects at micro- and macro-evolutionary scales. The detection of SVs is largely limited by the type and quality of sequencing technologies adopted, therefore genetic variability linked to SVs may remain undiscovered, especially in complex repetitive genomic regions. In this study, we used a combination of long-read and linked-read genome assemblies to investigate the occurrence of insertions and deletions across the chromosomes of 14 species of birds-of-paradise and two species of estrildid finches including highly repetitive W chromosomes. The species sampling encompasses most genera and representatives from all major clades of birds-of-paradise, allowing comparisons between individuals of the same species, genus, and family. We found the highest densities of SVs to be located on the microchromosomes and on the female-specific W chromosome. Genome assemblies of multiple individuals from the same species allowed us to compare the levels of genetic variability linked to SVs and single nucleotide polymorphisms (SNPs) on the W and other chromosomes. Our results demonstrate that the avian W chromosome harbours more genetic variability than previously thought and that its structure is shaped by the continuous accumulation and turnover of transposable element insertions, especially endogenous retroviruses.

## Introduction

Structural variants (SVs) are a heterogeneous category of DNA mutations that encompass changes in the copy number of a sequence (e.g., insertions, deletions, duplications, segmental duplications, transposable element insertions), changes in sequence orientation (e.g., inversions, translocations) and other changes in chromosome structure (e.g., chromosomal fusions/fissions, centromere shifts) (1, 2). It is thanks to continued advancements in genome sequencing and assembly that the relevance of SVs in evolution has begun to be appreciated (1, 3) in model and non-model organisms (4–7). Recent studies have found that SVs are common and may comprise the largest source of genetic variation within and between species (8–10). Evidence is also accumulating on the involvement of SVs in human diseases (11–13) and their role in the evolution of phenotypic traits (4, 14–17). SVs can have fitness effects through changes in gene expression but also by shaping the recombination landscape of chromosomes (18). Models of sex chromosome evolution also consider the role of inversions and insertions/deletions as key mutations for the stepwise vs. gradual recombination suppression between sex chromosomes (19–22). Given the frequent occurrence and importance of SVs, there is an increasing tendency to include the entire SV repertoire of a population or species into more complete and multi-individual reference genome assemblies called pan-genomes (23, 24). To proceed forward in determining the neutral, adaptive, and deleterious effects of SVs, as well as the type and strength of evolutionary forces acting on them, it is pivotal to broaden the characterisation of SVs to as many (non-model) organisms as possible.

The success in detecting and characterising SVs in genome assemblies is tightly linked to the type of sequencing techniques adopted, assembly contiguity, and intrinsic genomic features like the repetitive content (4, 25–27). Avian genomes present convenient genomic features that make them valuable for the study of SVs. These genomes, with few exceptions, are on average 10% repetitive and ∼1 Gb in size (28), in contrast to mammalian genomes which are usually ∼50% repetitive and ∼3 Gb in size (28). These aspects make avian genomes technically easier to assemble with respect to other classes of vertebrates (25, 27). However, birds exhibit a variety of chromosome types with different characteristics that pose different challenges for assembling complete genomes and to the full discovery of SVs. Avian chromosomes can be divided into macrochromosomes (>40 Mb), intermediate chromosomes (<40 Mb but >20 Mb), and microchromosomes (<20 Mb), the latter often challenging to assemble due to their high GC and G-quadruplex motif content (27, 29, 30). Microchromosomes also have a relatively higher density of genes and recombination rate than macrochromosomes (31, 32). This combination of high gene content and high recombination rate makes microchromosomes more permeable to selection while being less subject to linked selection and genetic drift in comparison to macrochromosomes (31). Therefore, avian genomes represent a good opportunity to investigate the occurrence and distribution of SVs in autosomes that evolve under a different balance of neutral and selective forces.

Sex in birds is genetically determined through a ZW sex chromosome system in which females are ZW heterogametic and males ZZ homogametic (33). The W chromosome of non-ratite birds is largely non-recombining except for a small pseudoautosomal region (34, 35), highly repetitive (27, 29, 34, 36, 37), and retains highly dosage-sensitive genes (i.e., under strong purifying selection (37– 39)). Theory predicts that the W chromosome should harbour lower genetic variability with respect to the Z chromosome and autosomes because of its smaller effective population size, recombination suppression, absence of male mutational input through spermatogenesis, and potential W sweeps due to tight linkage with the mitochondrial genome (40–44). Comprehensive studies on SNP variation in the coding regions of the W chromosomes of chicken (45) and flycatcher (46) support this prediction and found even less than expected genetic variability. The general lack of W-linked genetic variability led to the view of the W chromosome as a “sex-linked appendix” (46) with no particular functions besides carrying highly dosage-sensitive genes (37, 38). While the frequency of SNPs on the W has been investigated, W-linked genetic variation due to SVs has, to our knowledge, yet to be studied. The lack of knowledge regarding SVs on the W chromosome is tied to assembly issues since its repetitive content surpasses 70% in non-ratite birds (36), making it one of the most difficult-to-assemble avian chromosomes (27). The true levels of genetic variability on the W might have been hidden in past studies as part of the so-called genomic “dark matter” in contemporary genome assemblies (25, 27, 47). Assessing the level of W-linked variability with new long-read or multiplatform reference assemblies is necessary to better understand the magnitude of occurrence rate variation for different types of mutations (2) and to inform models of sex chromosome evolution (22).

Recent studies on birds have uncovered thousands of SVs on autosomes and the Z chromosome (4, 8), including SVs linked to reproductive behaviours (5) and speciation patterns (4, 48). However, none of these studies had female reference assemblies that would allow the identification of SVs carried by the W chromosome and were based on read mapping rather than assembly comparisons. In this study, we explored the occurrence, diversity, and distribution of SVs on autosomes and both sex chromosomes using linked-read draft assemblies from 14 species of birds-of-paradise (BOP; Paradisaeidae family), as well as long-read or multiplatform reference assemblies for two BOP species and two estrildid finch species. While the aforementioned studies focused on either very recent or very deep timescales of avian evolution, the present sampling of genome assemblies combined timescales which range from several individuals of the same species (the paradise crow *Lycocorax pyrrhopterus)*, the major branches of the BOP family, to species of another bird family (Estrildidae) which comprises the zebra finch *Taeniopygia guttata*. The comparison based on genome assemblies via pairwise whole-genome alignments against a high-quality reference assembly allowed for the investigation of regions that are difficult to survey with a read-based approach due to low and/or ambiguous mappability of such regions (49). Of the various types of structural variants, we focused on insertions and deletions because they are detectable with the genome assemblies available, while broader SVs (e.g., translocations, inversions) require additional data sources (e.g., optical maps or Hi-C (50)). Therefore, we were able to investigate the occurrence of insertions and deletions as well as the accumulation and turnover of transposable elements (TEs) on avian chromosomes including both sex chromosomes. In addition, thanks to the comparison of long-read assemblies for pairs of species in BOPs and Estrildidae, we were able to reconstruct the evolution of SVs (insertions and deletions) on autosomes and sex chromosomes in two bird families representative of the highly species-rich songbirds.

## Results

In this study, we investigated the evolution of insertions and deletions, as well as occurrences of interspersed repeats and tandem repeats at recent and deep evolutionary timescales, using genomic data ranging from individuals of the same species to species from different genera and families (**Figure 1**).

**Figure 1.**
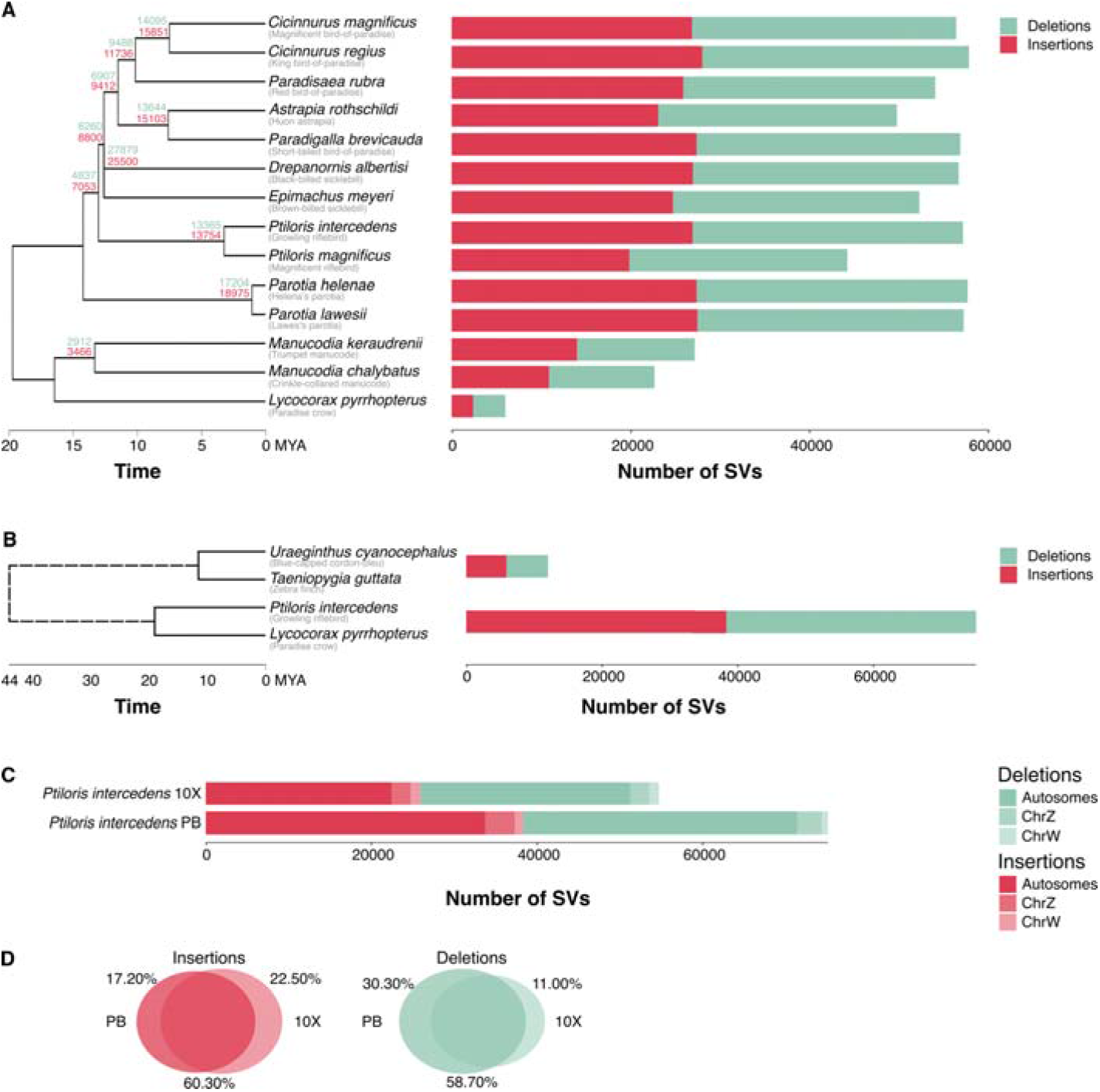
**A**) The genomes of birds-of-paradise and estrildid finches contain multitudes of SVs across evolutionary timescales and chromosome categories. The number of SVs identified with smartie-sv with respect to the reference assembly of *Lycocorax pyrrhopterus* is shown for 14 species of birds-of-paradise (BOPs) with 10X Genomics Chromium linked-read assemblies. The species tree follows the phylogeny from (53). The combined set of SVs called on the multiple *L. pyrrhopterus* individuals (F2-F4 and M1) is shown for the species. The numbers on top of the tree nodes indicate the number of polarised deletions (green) and insertions (red) relative to *L. pyrrhopterus* shared by all the species descending from that node. **B**) The number of SVs identified in species pairs with PacBio assemblies, i.e., between the two estrildid finch species *Uraeginthus cyanocephalus* against *Taeniopygia guttata* (reference) and between the two BOP species *Ptiloris intercedens* against *Lycocorax pyrrhopterus* (reference). **C**) Difference in the number of SVs identified in *P. intercedens* against *L. pyrrhopterus* using 10X Genomics linked-read (ptiInt10X) and PacBio long-read (ptiIntPB) assemblies. **D**) Percentage of SVs shared between the 10X Genomics and PacBio assembly comparisons of panel **C**.

We combined already existing genomic libraries and assemblies with newly produced sequencing libraries. We collected 1) the multiplatform female reference assembly of *Lycocorax pyrrhopterus* (27); 2) the 10X Genomics Chromium linked-read assemblies of three other females (F2-F4) and one male (M1) of *Lycocorax pyrrhopterus* (27, 36); 3) the multiplatform female reference assembly of *Taeniopygia guttata* (29); 4) re-sequencing Illumina libraries of one male museum sample each of *Cicinnurus regius, Cicinnurus magnificus, Epimachus meyeri, Manucodia chalybatus* and *Parotia helenae* (51). We generated new 10X Genomics Chromium linked-read libraries (i.e., short reads linked by unique barcodes) and assemblies of 1) females from 10 BOP species (of *Cicinnurus magnificus, Cicinnurus regius, Epimachus meyeri, Manucodia chalybatus, Manucodia keraudrenii, Parotia helenae, Parotia lawesi, Paradisaea rubra, Ptiloris intercedens* and *Ptiloris magnificus*, and 2) male from three BOP species (of *Astrapia rothschildi, Drepanornis albertisi* and *Paradigalla brevicauda*). In the newly produced 10X Genomics linked-read draft assemblies, the number of scaffolds ranged between 13,231 and 92,288 (26,236 on average) with a contig N50 between 27 kb and 187 kb (112 kb on average) and a scaffold N50 between 36 kb and 23 Mb (5 Mb on average). The most fragmented assembly (i.e., lowest contig N50) was that of *Ptiloris magnificus* while the most contiguous was that of *Drepanornis albertisi*. We also generated highly contiguous PacBio long-read assemblies for one female each of the BOP *Ptiloris intercedens* (880 contigs, contig N50 11.3 Mb) and the estrildid finch *Uraeginthus cyanocephalus* (942 contigs, contig N50 7.1 Mb). All assembly statistics are given in detail in **Table S1**.

### Structural changes within and between species

We investigated structural changes across chromosomes, with a particular focus on the sex chromosomes, by analysing the diversity and distribution of structural variants using all the genome assemblies available.

SV identification was carried out through smartie-sv (52) which compares contigs of target species to a reference assembly and can characterise insertions and deletions. The 10X Genomics draft assemblies of BOPs were each compared to the multiplatform reference of the paradise crow *Lycocorax pyrrhopterus*. In total 319,381 insertions and 355,035 deletions were identified (**Figure 1A**), which were on average 300 bp in size (minimum of 50 bp and maximum of 71,459 bp; **File S1, Figure S1**). The insertions affected 7.3 Mb on average in each genome (minimum of 250 kb in *Lycocorax pyrrhopterus* F4; maximum of 11.2 Mb in *Parotia lawesi*) while deletions affected 7.9 Mb on average in each genome (minimum of 578 kb in *Lycocorax pyrrhopterus* F4; maximum of 11.5 Mb in *Parotia lawesi*). The species presenting most SVs with respect to *L. pyrrhopterus* was *Cicinnurus regius* (**Files S1, Table S2**) while *Manucodia chalybatus* had the fewest such SVs. Very few SVs were called relative to the *L. pyrrhopterus* W chromosome in male individuals (27 on *Lycocorax pyrrhopterus*; 41 on *Astrapia rothschildi*; 61 on *Drepanornis albertisi*; 79 on *Paradigalla brevicauda*), these presumable misalignments between Z and W gametologs were not included in the tally and downstream analyses.

For *Ptiloris intercedens*, we ran smartie-sv against the *L. pyrrhopterus* reference with both the 10X Genomics (ptiInt10X) and the PacBio (ptiIntPB) assembly, obtaining different numbers of SVs (**Figure 1B** and **1C**). We identified 73,289 SVs (35,849 deletions and 37,440 insertions) using ptiIntPB and 54,055 SVs using ptiInt10X (28,436 deletions and 25,619 insertions). While with ptiIntPB we identified 19,819 more SVs on autosomes and the Z chromosome, 585 fewer SVs were identified on the W chromosome with respect to ptiInt10X (**Table S3**). The mean size of SVs in ptiInt10X was 397 bp (maximum of 37 kb), whereas in ptiIntPB the mean size was 466 bp (maximum of 69 kb: **Figure S1**). By intersecting the SV sets from ptiInt10X and ptiIntPB, we found that 58.70% of the deletions and 60.30% of the insertions were shared by both types of assemblies (**Figure 1D, Table S3**). Finally, the density distributions of SVs across chromosome categories were similar when calculated using the two assembly types (**Figure 2B**), except for microchromosomes where ptiIntPB showed higher between-chromosome variability of densities.

**Figure 2.**
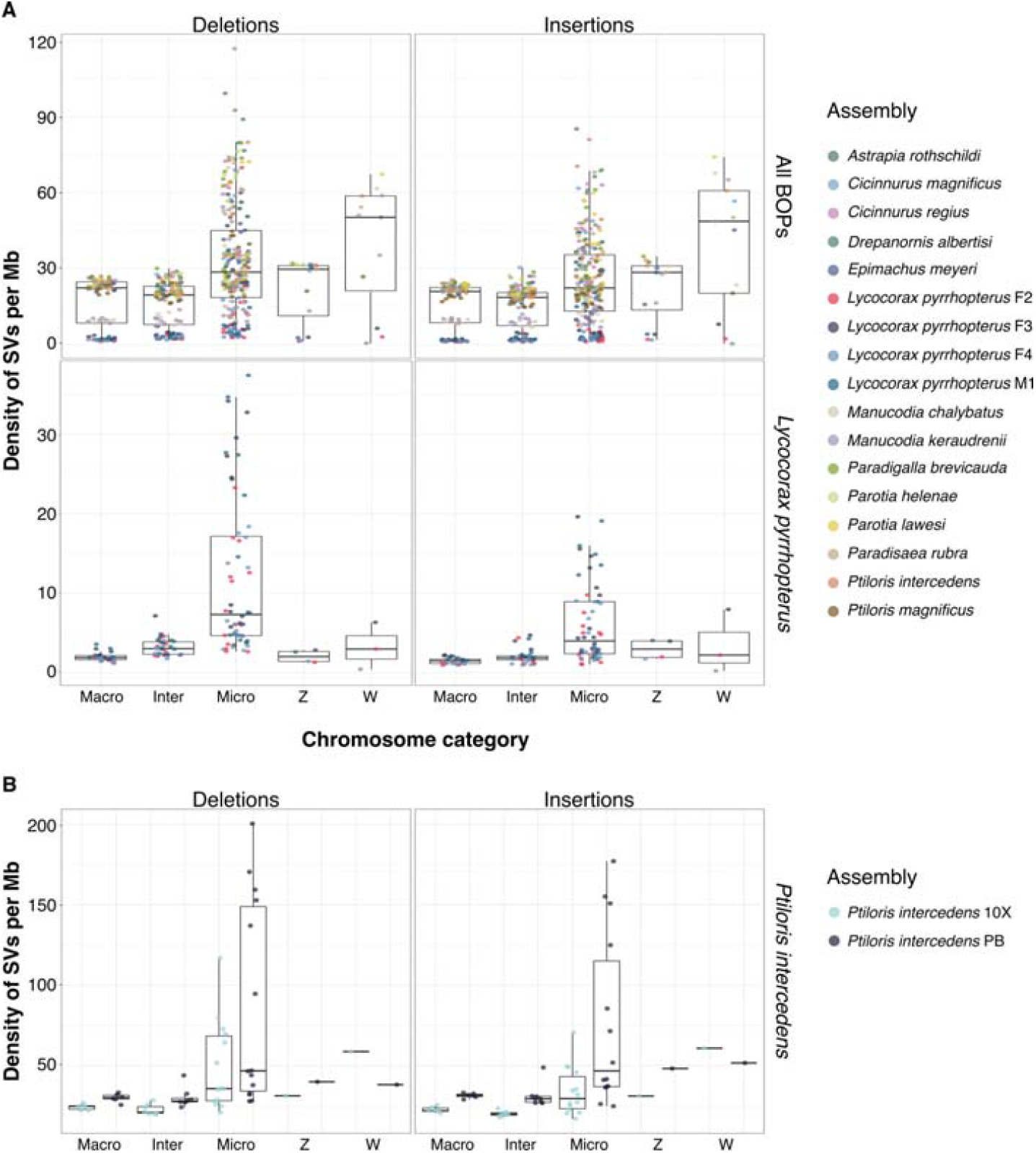
**A**) Density of SVs per megabase (Mb) identified with respect to the reference assembly of *Lycocorax pyrrhopterus* divided per chromosome categories. Density of insertions and deletions identified by smartie-sv in all the BOP species (**top panel**) and in the four individuals (three females and one male) of *Lycocorax pyrrhopterus* (**bottom panel**). **B**) Density of structural variants per Mb across chromosome categories. Density of deletions (**left panel**) and insertion (**right panel**) identified in *P. intercedens* against *L. pyrrhopterus* using both the 10X Genomics and PacBio assemblies of *P. intercedens*. Macro: macrochromosomes (>40 Mb); Inter: intermediate chromosomes (>20 Mb and <40 Mb); Micro: microchromosomes (<20 Mb).

In general, microchromosomes showed more insertions and deletions relative to their size than other chromosomes (**Figure 2A**). This applied both to the comparisons between BOP species (**Figure 2A top panel**) and between *L. pyrrhopterus* individuals (**Figure 2A bottom panel**), with the exception of the W chromosome. While the density of SVs on the W chromosome was within the range of values of the other chromosomes when other *L. pyrrhopterus* individuals were taken into consideration, it was higher when other species were compared. Not considering the *L. pyrrhopterus* samples, the species with most SVs on the W (1,324 deletions and 1,458 insertions) was *Parotia lawesi* and the species with fewest was *Manucodia chalybatus* (417 deletions and 464 insertions; **File S1**).

The analyses of female PacBio assembly of *Uraeginthus cyanocephalus* (ura-CyaPB) against multiplatform reference assembly of *Taeniopygia guttata* revealed 13,362 SVs, of which 6,772 were deletions (**Figure 1B**) that affected 2.23 Mb and 6,590 were insertions that affected 2.15 Mb. No W-linked SVs identified in this species comparison passed the filtering step.

Then, we characterised which SVs were shared between BOP species at the different nodes in the phylogeny. To do that, we first overlapped the SV called against *L. pyrrhopterus* from species belonging to the terminal nodes of the phylogeny and then walked node-by-node backwards across the phylogeny (**Figure 1A**). Deletions with a reciprocal overlap of at least 70% and insertions within 50 bp from one another were considered shared. We found that closely related species (e.g., species of the same genus) shared about 50% of their SVs and were able to anchor thousands of SVs to deeper branches of the phylogeny.

Afterwards, we compared the overall levels of genetic variability linked to SVs and SNPs across chromosomes. We did so using multiple individuals of *L. pyrrhopterus* and calculating four statistics; SV density per window, SNP density per window, nucleotide diversity (π) using pixy (54), and SV diversity (see **Methods**). The density and diversity values calculated for windows of 100 kb for the sex chromosomes and the autosomes 1, 5, and 18 in order to encompass different chromosome sizes are depicted in **Figure 3**. SV density was low for every analysed chromosome, with chromosome 1 showing the lowest SV density (0.000008) and chromosomes 18 and W showing the highest SV densities (0.000026 and 0.000019 respectively; **Table S3**). Regarding SNP density, the autosomes showed the highest values, and the Z chromosome showed half the mean density of the autosomes (0.0046 to 0.0050 vs. 0.0018). The W chromosome had a mean SNP density of 0.0002 (**Table S4**). In general, SNP density was significantly higher on all the autosomes with respect to the Z and W (Kolmogorov-Smirnoff test, p-values <2.2e-16). Furthermore, mean nucleotide diversity (π) between autosomes was very similar (ranging between 0.002 and 0.0025), while π was half of this on the Z chromosome (0.001). However, the W chromosome showed a π value of 0.0018, i.e., similar to the autosomes (**Table S5**). Regarding SV diversity, mean values were 3.69e-07 (chr1), 3.46e-07 (chr5), 1.03e-06 (chr18) on autosomes, 2.95e-07 on the W, and were lowest on the Z with 1.08e-07. The distributions of SV diversity values on the autosomes were not statistically different from the Z and W (Wilcox-test, lowest p-value: 0.1) except for chromosome 18 (highest p-value: 5.2e-06).

**Figure 3.**
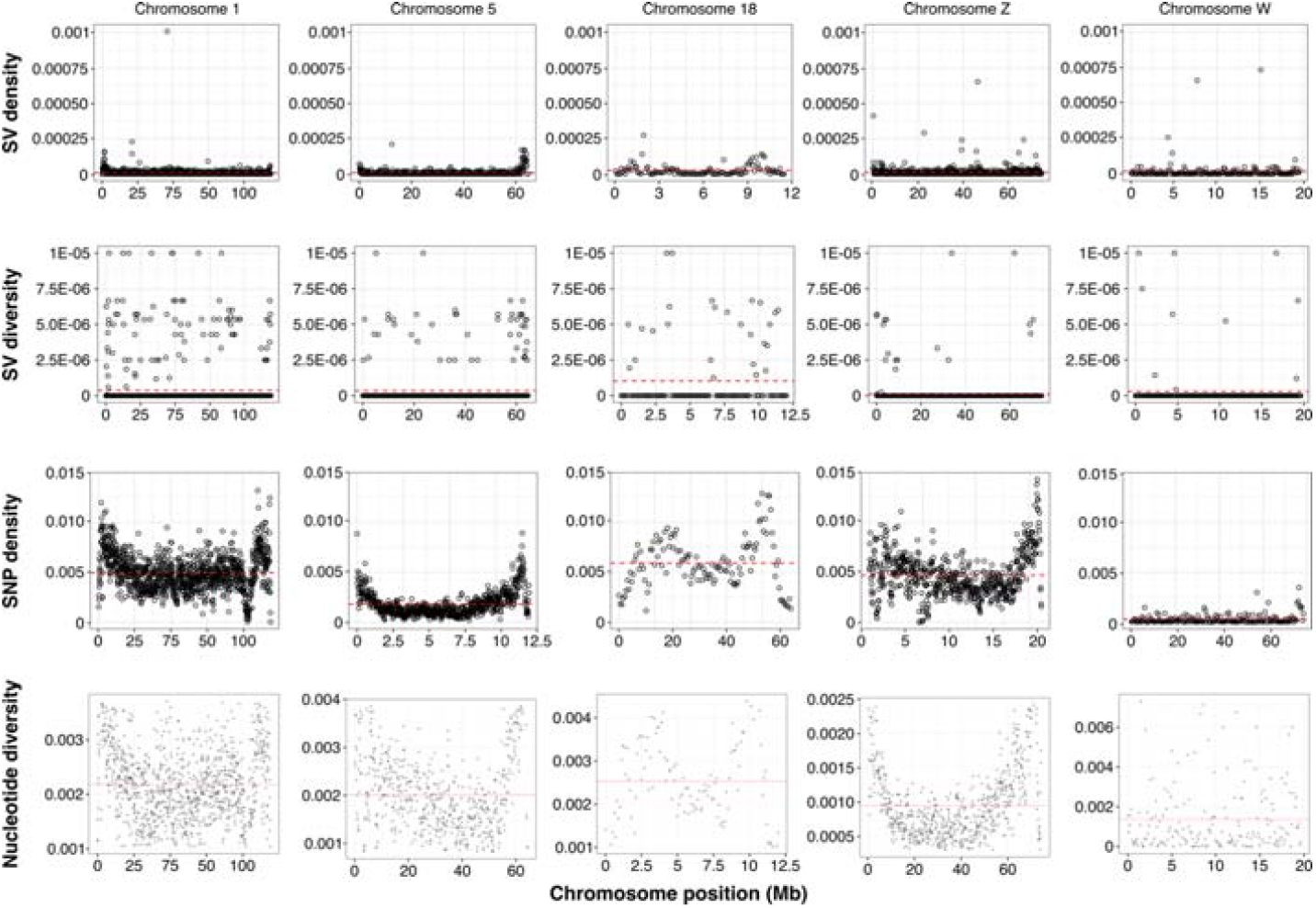
The landscape of density and diversity of structural variants and SNPs across sex chromosomes and selected autosomes of *Lycocorax pyrrhopterus*. Values are plotted per window of 100 kb. The mean values for each statistic are shown as dashed red lines.

### SVs caused by repeats and other mechanisms

To understand which mechanisms may have caused the observed SVs at different timescales, we investigated two plausible explanations for insertion/deletion occurrence: 1) TE insertions and other repetitive elements, and 2) recombination and DNA repair mechanisms.

First, we investigated whether these SVs were the result of TE insertions. TEs are DNA sequences with their own means of mobility and insertion across the host genome, therefore their insertions are a specific type of insertion/deletion SVs (2). For the SV annotation and repeat annotation to be considered overlapping, we required the SV position to be masked for at least 70% of their length by one or more repeats. When considering SVs identified between *Lycocorax pyrrhopterus* individuals, 35-40% of SVs overlapped with repeats. Only 10-14% of all the SVs overlapped with TEs, of which 6-10% were endogenous retroviruses (ERVs), a group of long terminal repeat (LTR) retrotransposons, and 2-4% were CR1 LINE retrotransposons. Another 20% of SVs overlapped with simple repeats and complex repetitive regions, i.e., regions occupied by multiple types of repeats. The percentages of SVs co-occurring with repetitive elements were ∼60% when genomes of other BOP species were compared to *Lycocorax pyrrhopterus*. In these comparisons, 27-41% of SVs co-localised with ERVs, 5% with CR1 LINEs, 8-10% with simple repeats, and 5-6% with complex repetitive regions (**Table S2**). When comparing the SVs called by ptiIntPB and ptiInt10X, we observed that in ptiIntPB more SVs were masked as satellite DNA and simple repeats, and fewer SVs were complex repetitive regions. The respective percentages of SVs overlapping with TEs were 32% and 44% in ptiIntPB and ptiInt10X (CR1 LINEs 2% and 5%, ERVs 30% and 39%, SINEs 8% and 20%; **Table S2**).

SVs commonly arise as by-products of meiotic recombination or DNA repair events (55). To check whether those SVs that cannot be explained as a result of insertions/deletions of TEs may be linked to recombination or DNA repair events, we analysed the degree of homology shared by the flanking regions of the SVs not overlapping with repetitive elements, using approaches from human SV studies (52, 55). A total 3-4% of the insertions and deletions identified among the *L. pyrrhopterus* samples could be explained by events of non-allelic homologous recombination (NAHR; >200 bp homology between flanks), homology-directed repair (HDR; >50 bp homology between flanks) and to microhomology-mediated end joining or microhomology-mediated break-induced replication (**Table S4**). The percentages were 0.5-1.5% when analysing the other BOP species (**Table S4**).

### Interspersed repeat evolution

As tested above, structural variants can originate from events of NAHR (55, 56) between (near-)identical sequences, for example between or within individual TE copies. The rate of NAHR events is expected to follow the rate of homologous recombination (57). Since avian genomes exhibit chromosomes with very different recombination rates (31, 58–60), we tested whether the rate of NAHR events follow these expectations or whether there are chromosomes in which this mechanism is more/less accentuated. A proxy for the occurrence of NAHR events is the ratio between solo and full-length LTR retrotransposons (61). LTR retrotransposons can be found in the genome in their full length (characterised by an internal protein-coding portion flanked by two long terminal repeats, the LTRs, at the extremities, which are identical upon insertion) and in their solo LTR forms (21, 62). The latter form is the result of an NAHR event between the two LTRs, removing one of the two LTRs and the internal portion (21, 62). The solo-to-full-length ratio was estimated as a proxy for the rate of conversion of full-length LTRs in solos, which we consider as a proxy for the rate of NAHR. A high ratio should correspond to a fast full-length to solo conversion. For this analysis, we used only the multiplatform reference assemblies of *L. pyrrhopterus* and *T. guttata* as they have chromosome models assembled (27, 29).

The ratios on the autosomes mostly ranged between 10 and 100, with a few microchromosomes (19, 20, 24, 26) where ratio values were more similar to the Z chromosome (ratio of 7), despite the expectation that autosome sizes inversely correlate with recombination rates (31, 58–60). Chromosomes 28 and 33 appeared to lack full-length LTR retrotransposons despite the presence of solo LTRs, therefore no ratio was estimated for these chromosomes (**Figure 4**). Given the very low (3-4) expected number of full-length LTR retrotransposons on such small microchromosomes, their absence might be stochastic or reflect limitations of the assembly and detection tools, rather than reflect the biology of these chromosomes. Finally, the solo-to-full-length ratios were the lowest on the male-recombining Z chromosome and on the non-recombining female-specific W chromosome (**Figure 4, File S2**), the latter being the most highly enriched for full-length LTR retrotransposons.

**Figure 4.**
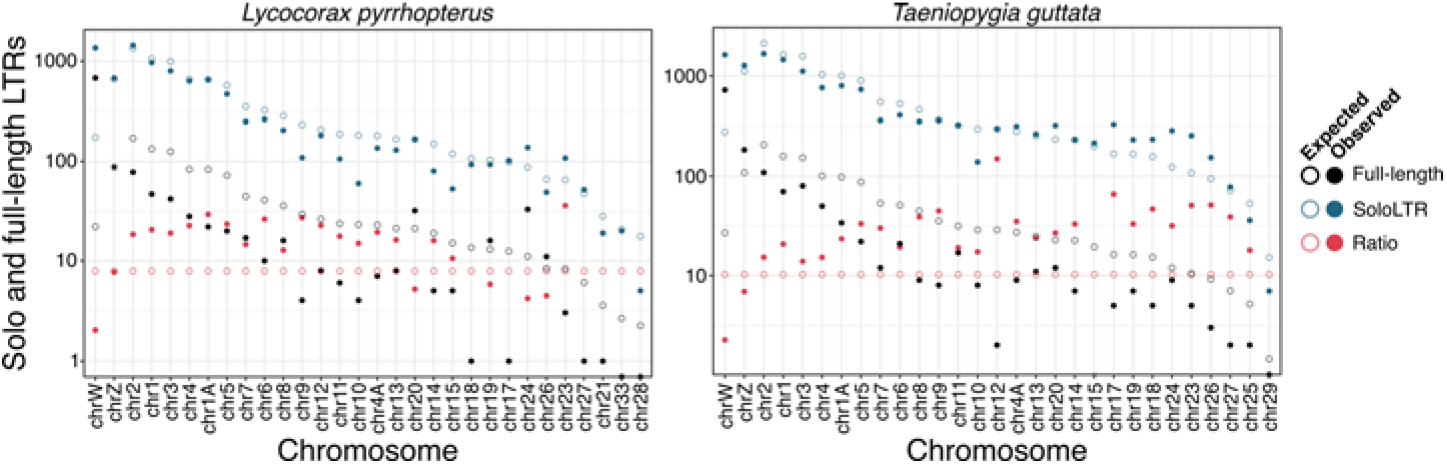
The W chromosome and microchromosomes are highly enriched for full-length LTR retrotransposons and solo LTRs, respectively. The number of expected and observed solo and full-length LTR retrotransposons and their ratios are shown for each chromosome of *Lycocorax pyrrhopterus* and *Taeniopygia guttata* reference genomes. A high ratio indicates that more solo LTRs are present than full-length LTRs, while a low ratio indicates the opposite. Each X-axis first lists the non-recombining W chromosome and the male-recombining Z chromosome, followed by autosomes sorted inversely by chromosome size as proxy for their increasing recombination rate. The Y-axis is given in log10 scale.

To further understand if TEs evolved differently on the W chromosome with respect to the Z and autosomes, we compared the relative age of W-linked TEs to TEs on the other chromosomes. The relative age of a TE copy is commonly inferred by the number of mutations accumulated relative to a consensus sequence (63). The consensus sequence of a TE is the sequence approximation of the ancestral sequence that gave rise to the TE copies of each TE subfamily (64). Assuming neutrality, the more substitutions are present on the annotated TE insertion with respect to the consensus sequence, the older the insertion is. Therefore, we tested whether the W chromosome accumulated new TE insertions faster than the other chromosomes and whether TE insertions on the W aged (mutated) faster than those on the Z and autosomes.

Given the lack of recombination and thus low effective population size of the W, we expect this chromosome to accumulate new TE insertions faster than the other chromosomes, as well as to have a faster TE turnover. Therefore, we also expect that the continuous accumulation of new insertions should drive the average age (mean divergence) of active TE subfamilies down. While insertions of active TE subfamilies keep accumulating, the past insertions of inactive subfamilies are not supplemented anymore by new copies and thus the mean divergence from consensus increases. To test this expectation, we calculated the weighted mean of the divergence from consensus of the TE insertions on the different chromosome categories (autosomes, Z, and W) and compared the distribution of these means in young and old TE subfamilies using the Kolmogorov-Smirnoff (K-S) test (**File S7**). A TE subfamily was considered young when it presented insertions with a divergence of 5% or less. On the other hand, a TE subfamily was considered old when all of its insertions showed a divergence >10%. In *Lycocorax pyrrhopterus*, when 312 young subfamilies were compared, subfamilies on the W looked significantly younger with respect to the autosomes (K-S test; p-value 0.0002) and to the Z chromosome (K-S test; p-value 7.922e-07). Similarly, also subfamilies on the Z looked younger than on the autosomes (K-S test; p-value 0.0488). On the other hand, when the 626 old subfamilies were considered, they looked older on the W chromosome with respect to the autosomes (K-S test; p-value 7.72e-05) but not to the Z chromosome (K-S test; p-value 0.07277); likewise, the old subfamilies on the Z looked older than on the autosomes (p-value: 0.002365). In *Taeniopygia gutatta*, we found 437 young subfamilies that looked younger on the W chromosome with respect to the Z (K-S test; p-value 0.0007) but not to the autosomes (K-S test; p-value 0.038). The 610 TE subfamilies labelled as old were found to look older on the W with respect to the autosomes (K-S test; p-value 0.0001) and on the Z with respect to the autosomes (K-S test; p-value 0.0001). The detailed results of the Kolmogorov-Smirnoff test (including the D scores) are shown in **File S6**.

## Discussion and conclusions

Structural variants reperesent an important source of genetic variation and are thought to play a crucial role in the evolutionary process (1–4, 9, 10). Yet to understand it is how this source of variation contributes to the evolutionary dynamics at both macro and microevolutionary levels, it is therefore key to document their importance in genomes across a diverse range of organisms. Here, we explored the occurrence of SVs and repetitive elements in 14 species of BOPs and two species of estrildid finches using both (linked) short-read and long-read data (**Figure 1A** and **B**). Using a multiplatform reference genome assembly we called SVs using both linked-read draft assemblies and high-quality long-read assemblies. We found SVs occurring on all chromosomes (**Figure 2A**), including the non-recombining female-specific W chromosome which, to our knowledge, had previously not been characterised for SVs in any bird. Since avian karyotypes present macrochromosomes, intermediate chromosomes, and microchromosomes (32), we divided the resulting SVs into these chromosome categories and found that microchromosomes show a higher density of SVs with respect to the other categories both at inter-species and intra-species levels (**Figure 2A**). In addition, the W chromosome also exhibits a high density of SVs especially at the interspecific level. We propose that this high density of SVs can be linked to sequence composition of this chromosome. The W chromosome in Neognathae birds is highly heteromorphic (34) and repetitive (27, 36), featuring most of the full-length TEs of the genome (especially potentially active ERVs) as well as most of the young insertions with a low divergence from consensus (36). Indeed, our results further strengthen the observations that the W chromosome accumulates new insertions more rapidly than other chromosomes, highlighted by a lower average divergence of recently mobilised TE subfamilies on the W with respect to the Z and the autosomes (**File S3**). Conversely, we also noted that old TE subfamilies show a higher average divergence on the W. These two divergence patterns suggest that, while W-linked TE insertions experience a higher rate of substitution with respect to the other chromosomes as expected from the lower effective population size and higher effect of genetic drift on the W (42), the continuous accumulation of recent insertions lowers their mean divergence from consensus. Once the TE subfamilies become inactive and no new insertions accumulate on the W anymore, the W-linked insertions diverge from the consensus faster than on other chromosomes.

The tendency of the W to accumulate new insertions in great quantity provides, theoretically, the optimal homogeneous sequence substrate to trigger events of non-allelic homologous recombination (NAHR), a form of ectopic recombination commonly associated with structural rearrangements (55). Assuming that the rate of NAHR is generally correlated with the rate of homologous recombination, we estimated the solo-to-full-length ratio of LTR retrotransposons. By calculating this ratio in BOPs and estrildid finches (**Figure 4**), we found that more NAHR is generally associated with a higher recombination rate of chromosomes as expected (57). Indeed, the solo-to-full-length ratio is lowest on the W chromosome, which is non-recombining outside a small pseudoautosomal region, and in the Z chromosome which fully recombines only in males, i.e., half of the population. The patterns on the microchromosomes are less clear though. Microchromosomes have a higher recombination rate than macrochromosomes and intermediate chromosomes (31, 58–60), so a higher solo-to-full-length ratio is expected with decreasing chromosome size. While *T. guttata* shows this expected pattern, most microchromosomes in *L. pyrrhopterus* do not show such a pattern(rather ratios similar or lower than macrochromosomes). While the discordant patterns between the two birds can be biological, it must also be considered that limitations in the assembly and/or LTR detection tools could be a partial cause of such discordance especially on the GC-rich microchromosomes. Our results suggest that the expected number of full-length LTRs in the smallest microchromosomes is very low (between 1 and 10) and together with the fact that some microchromosomes are particularly difficult to assemble correctly (65), it is probable that those sequences could be missing or misassembled. From the solo-to-full-length ratio results, we gather that NAHR occurs also on the non-recombining W chromosome and that the rate of solo LTR formation is faster than the rate of insertion of new full-length LTR retrotransposons. However, it must be noted that when we investigated what types of mechanisms could underlie the SVs detected in the pairwise comparison of genome assemblies, we found only 1-4% of the SVs to show evidence of NAHR. Alternatively, this result might be a by-product of the draft quality of the linked-read assemblies used to call the SVs and thus might reflect uncertainty in the detection of precise SV breakpoints.

Our results also suggest that the choice of sequencing technology to use for SV calling is essential to get reliable calls (see also (4)) and to be able to compare SVs from different species. Given our genome assemblies were based on two types of sequencing data (linked and long reads), we explored the effects of using different genomic data for the same species on SV calling. When comparing the SVs called by using a 10X Genomics linked-read assembly and a PacBio long-read assembly of *P. intercedens* (with *L. pyrrhopterus* as reference), we found the two sets of SVs to share most of the calls (58-60% of the SVs were called by both assemblies). Whereas part of the non-shared SVs can be false positives, some of these SVs are likely detectable only with the long-read assembly where these regions are assembled.

After investigating SVs between different species, we focused on SVs occurring between individuals of the same species using the *L. pyrrhopterus* samples and compared the SV-linked diversity across chromosomes to the SNP-linked diversity. We investigated the genetic diversity of the sampled *L. pyrrhopterus* population through the SNP and SV density, the nucleotide diversity π, and an adapted version of π for SVs. The SNP density on the W chromosome was low with respect to the Z and autosomes, but its nucleotide diversity similar to the other chromosomes (**Figure 3**), which contrasts with previous results in chicken and flycatcher (45, 46). We suspect that these unexpected π values can be linked to the fact that previous studies focused only on coding regions and therefore had underestimated π. The SV density and diversity showed values whose distributions were similar between the W and other chromosomes.

We consider our estimates of SV density and diversity to be very conservative since we were comparing draft assemblies to long-read assemblies with strict filters. We expect not all regions of a genome to be equally scorable for SVs, especially highly repetitive ones like the W. Now with a high-quality but not gap-free assembly of the W, the situation looks almost paradoxical where assembly completeness entails scorability issues. While the W assembly is getting more and more complete in multiplatform assemblies, we suspect that the now assembled repeats are in turn the least scorable regions for the detection of SVs. Even with these conservative estimates regarding SV presence, the W here showed values of SV density and diversity similar to the other chromosomes, i.e., the true values are probably much higher.

We hypothesise that a highly repetitive W chromosome with a continuous accumulation of potentially active LTR retrotransposons (36) and an SV diversity similar to or higher than the other chromosomes, can lead to the frequent reshaping of the heterochromatic landscape of the female-specific W chromosome itself as well as of the entire genome. Studies on *Drosophila melanogaster* highlighted that SVs involving repetitive regions on the male-specific Y chromosome can have epistatic effects on the expression of genes across the entire genome (66– 69). The dynamic structural variability of the W, therefore, may have epistatic and fitness effects as well as the potential to solve genetic conflicts by modulating gene expression. We predict that the thorough investigation of additional types of SVs on ever-improving W chromosome assemblies will continue to provide a better understanding of sex chromosome evolution and of the different mutational forces shaping (avian) genomes in general.

## Methods

### Samples and sequencing libraries

For this study, we used previously published as well as newly produced genomic data. We retrieved the high-quality chromosome-level genome assembly of *Lycocorax pyrrhopterus* (27) and 10X Genomics Chromium linked-read libraries (i.e., short reads linked by unique barcodes) and assemblies for three female and one male of the same species (27, 36). The DNA for one female individual (pectoral muscle) each of *Cicinnurus regius, Cicinnurus magnificus, Paradisaea rubra, Epimachus meyeri, Ptiloris intercedens, Ptiloris magnificus, Parotia helenae, Parotia lawesi, Manucodia keraudrenii, Manucodia chalybatus* and one male individual (pectoral muscle) each of *Astrapia rothschildi, Drepanornis albertisi, Paradigalla brevicauda* were extracted with the Kingfisher Duo robot using the KingFisher Cell and Tissue DNA Kit following the manufacturer’s recommendation and eluted in 100 μl elution buffer. For *Uraeginthus cyanocephalus*, high molecular weight DNA was extracted from female blood at SciLifeLab Uppsala using the Circulomics Nanobind kit. For *Ptiloris magnificus*, sampling in Papua was conducted according to relevant research and ethical guidelines by the government of the Republic of Indonesia and under research permits issued by RISTEK (Indonesia) (304/SIP/FRP/SM/X/2014) and relevant Indonesian government collecting permits.

10X Genomics Chromium libraries were generated from these samples and sequenced at SciLifeLab Stockholm, either on an Illumina HiSeq X instrument or an Illumina NovaSeq 6000 instrument. Low-coverage (5-8X) Illumina libraries for one male each of *Cicinnurus regius, Cicinnurus magnificus, Parotia helenae, Manucodia chalybatus* and *Epimachus meyeri* were retrieved from (51). In addition, we newly generated a PacBio long-read library of the same *Ptiloris intercedens* female sample (104 Gb total data, 10 kb read N50, 15 SMRT cells, Sequel II) used for generating the 10X Genomics Chromium linked-read library, and a PacBio HiFi long-read library for a female sample of *Uraeginthus cyanocephalus* (58 Gb total data, 20 kb mean read length, two SMRT cells, Sequel II HiFi mode). PacBio library generation and sequencing was done at SciLifeLab Uppsala.

### Genome assembly generation

The 10X Genomics Chromium libraries for the species listed above were assembled *de novo* with the dedicated assembler Supernova2 (70) and the pseudohaploid assembly versions were used for downstream analyses. The PacBio library obtained for *Ptiloris intercedens* was assembled into primary contigs by the diploid-aware assembler Falcon-unzip (71). The primary contigs were polished with a round of Arrow (https://github.com/PacificBiosciences/GenomicConsensus), two rounds of Pilon (72) and scaffolded with ARCS (73) and LINKS (74) using the 10X Genomics Chromium library for the same individual following the methods and parameters in (27). The PacBio HiFi library for *Uraeginthus cyanocephalus* was assembled using IPA (https://github.com/PacificBiosciences/pbipa). Uncollapsed haplotypes were removed from the assembly of both *P. intercedens* and *U. cyanocephalus* with Purge Haplotigs (75).

### Structural variant calling

We used smartie-sv (52) to call structural variants present in the BOP linked-read draft assemblies with respect to the *L. pyrrhopterus* reference assembly, the *P. intercedens* long-read assembly with respect to the *L. pyrrhopterus* reference assembly, and the *U. cyanocephalus* long-read assembly with respect to the *T. guttata* reference assembly. Smartie-sv takes contigs as input, therefore we split the assemblies back into contigs when necessary. Only the SVs that 1) were located within chromosome models (i.e., SVs within unknown chromosomes, unplaced scaffolds and contigs were discarded), 2) contained less than 10% of N nucleotides in their sequences, and 3) showed an alignment identity higher than 90% were used for downstream analyses. To find and merge overlapping SVs among the multiple individuals of *L. pyrrhopterus*, we concatenated the smartie-sv outputs and merged the coordinates of the SVs using BEDTools merge (76). The W-linked SVs identified using the male *L. pyrrhopterus* assembly were used as a blacklist of false positives. These false positives, as well as all the SVs overlapping with them, were discarded from downstream analyses.

We calculated the density of SVs per megabase for different chromosome categories by dividing the chromosomes into macrochromosomes (>40 Mb), intermediate chromosomes (20-40 Mb), microchromosomes (<20 Mb), and Z and W. Finally, to identify which structural variants overlapped with repetitive elements, we intersected the coordinates of the SVs with the RepeatMasker output using BEDtools intersect (76) requiring a reciprocal 70% of overlap (-f 0.7 -r).

Finally, using the phylogeny from (53), we identified which SVs were shared between BOP species at the different nodes in the phylogeny. We walked node-by-node backwards across the phylogeny and overlapped the deletion and insertion coordinates between species. Deletions with a reciprocal overlap of at least 70% and insertions within 50 bp from one another were considered shared. The number of shared SVs at the deepest nodes of the phylogeny were not reported because, given the species sampling, those SVs cannot be polarised.

### SNP and SV diversity

To investigate the diversity of SVs and SNPs at the population level, we calculated nucleotide diversity (pi) and structural variant diversity as well as SNP and SV densities. First, we ran pixy (54) to calculate the nucleotide diversity within the *L. pyrrhopterus* population. Pixy is a tool designed to minimise biases in the calculation due to missing genotypes or sites. To use pixy, we first mapped the 10X Genomics libraries of four *L. pyrrhopterus* females and one male to the *L. pyrrhopterus* reference genome assembly with Longranger. The obtained alignment files were combined with bcftools (77) to generate a comprehensive VCF file that included the invariant sites following the suggested code given by pixy (https://pixy.readthedocs.io/en/latest/index.html). The VCF file was then used to calculate the nucleotide diversity per window of 100 kb. Second, the same VCF file was filtered for SNPs with a quality phred score higher than 30 and the number of SNPs per windows (100 kb) was calculated by using BEDTools intersect. Before estimating the SNP density, the size of the windows was corrected for the presence of N nucleotides. Similarly, we calculated the density of SVs by intersecting the SV and window coordinates, and correcting the window sizes for the presence of N nucleotides as they constitute missing sites.

Finally, we estimated the SV diversity in the population by genotyping the SVs with Paragraph (78) and custom calculations adapting the approach implemented in pixy for nucleotide diversity. The calculation of nucleotide diversity using genotyped SVs is a way to estimate the levels of polymorphisms in the population from an SV point of view. To run Paragraph, we used the alignment files produced with Longranger and SV coordinates converted into a VCF file as inputs. The resulting genotype VCF file was parsed and turned into a table of haplotypes where all the genotypes marked as “.” in the GT field or marked as “CONFLICT”, “NO_VALID_GT”, “NO_READS”, “UNMATCHED” or “BP_DEPTH” were considered as missing genotypes. After filtering, we calculated the nucleotide diversity on this set of haplotypes while fixing the number of comparisons, taking the missing genotypes into account and adjusting the window sizes for the presence of N nucleotides.

### Solo/full-length LTR ratio and repetitive element turnover

We used available full-length LTR annotations for *L. pyrrhopterus* and *T. guttata* from (36) and carried out the annotation of solo LTRs using the findSoloLTRs pipeline developed for this study (https://github.com/ValentinaBoP/Wevolution). The findSoloLTRs pipeline took as primary input the RepeatMasker output (.out) file and the genome assembly of interest. The RepeatMasker output file were filtered for retaining only hits belonging to the long terminal repeats of the ERVs that spanned the entire length of the matching consensus sequence (5 bp upstream and 5 bp downstream were given as threshold of completeness). All the hits belonging to the internal portion of the LTR elements, to elements labelled as incomplete (.inc) or “LTR?”, and fragmented hits were discarded. To find target site duplications (TSDs) that are the hallmark of an actual transposition event, the proximal 10 bp upstream and 10 bp downstream to the complete hits were extracted from the assembly and aligned to one another using Blast (79). The common length of TSDs for LTR retrotransposons spans from 4 to 6 bp (80), therefore we used a word size of 4 in BLAST to accommodate for such microhomologies. All TSDs that appeared to be longer than 6 bp were discarded at the end of the pipeline.

After retrieving the coordinates of both solo and full-length LTRs, we calculated the solo-to-full-length ratio (62) for each chromosome. This ratio is a proxy for the speed of the conversion of full-length elements into solo LTRs and of the rate of non-allelic homologous recombination (NAHR). We also calculated the expected number of full-length and solo LTRs as well as the expected ratio by assuming a homogeneous density of these elements across the chromosomes.

To investigate how fast TE insertions mutate on the W, Z, and autosomes with respect to their sequence at time point of insertion (approximated by the consensus sequence), we compared the genetic divergence of TE from their consensus sequences on the different types of chromosomes using the RepeatMasker annotations for *L. pyrrhopterus* and *T. guttata*. If TEs mutate at the same rate on all chromosomes, then we expect the subfamilies to have a similar divergence from consensus in all the chromosomes. To test this, we identified young and old sub-families and compared their average divergence from consensus. First, we calculated the weighted mean genetic divergence for each TE subfamily annotated by RepeatMasker on each chromosome category (autosomes, Z, and W). Second, we established a criterion to distinguish young and old TE subfamilies: We considered young those TE subfamilies which presented insertions with a divergence of 5% or less; similarly, we considered old those subfamilies which present zero insertions at 10% divergence and below. Then, we compared the distributions of the weighted means of young and old TE subfamilies between the different chromosomes applying the Kolmogorov-Smirnoff non-parametric (one-sided) test using the ks.test function in R (81).

## Acknowledgements

We thank Antje Bakker for technical assistance, David Witkowski for animal care, and the Suh laboratory, Johannesson laboratory, Matthias Weissensteiner, Reto Burri, and Marco Ricci for helpful discussions. This work was supported by the Kungliga Fysiografiska Sällskapet i Lund (Nilsson-Ehle Donation to V.P.), the Swedish Research Council Vetenskapsrådet (2016-05139 to A.S.; 621-2014-5113 and 2019-03900 to M.I.), and the Swedish Research Council Formas (2017-01597 to A.S.). Samples for this study were kindly provided by Museums Victoria in Melbourne (MV), the Australian National Wildlife Collection in Canberra (ANWC), the Peabody Museum of Natural History and Yale University Museum in New Haven (YPM), the Swedish Museum ofs Natural History in Stockholm (NRM), the Natural History Museum of Denmark in Copenhagen (ZMUC), and the University of Kansas Biodiversity Institute (KU). We are particularly grateful to Joanna Sumner, Leo Joseph, Robert Moyle, Ulf Johansson, and Knud Jønsson for the assistance with this. Fieldwork in West Papua, Indonesia, was supported by the Lengguru Project (www.lengguru.org) conducted by the French Institut de Recherche pour le Développement (IRD), the Indonesian Institute of Sciences (LIPI) with the Research Center for Biology (RCB) and the Research Center for Oceanography (RCO), the University of Papua (UNIPA), the University of Cendrawasih (UNCEN), the University of Musamus (UNMUS), and the Polytechnic KP Sorong with corporate sponsorship from COLAS and TIPCO groups, Veolia Water and the Total Foundation. Long-read sequencing data was generated by Olga Vinnere-Pettersson, Susanne Hellstedt Kerje, Ignas Bunikis, Mai-Britt Mosbech, Sara Olofsson, and Pernilla Quarfordt at the National Genomics Infrastructure in Uppsala (Uppsala Genome Center), and linked-read sequencing data by Max Käller, Mattias Ormestad, Elísabet Einarsdóttir, Remi-André Olsen, Joel Gruselius, Fanny Taborsak-Lines, and Franziska Bonath at the National Genomics Infrastructure in Stockholm. Both facilities are funded by Science for Life Laboratory, the Knut and Alice Wallenberg Foundation and the Swedish Research Council. Computations were performed on resources provided by the Swedish National Infrastructure for Computing (SNIC) through Uppsala Multidisciplinary Center for Advanced Computational Science (UPPMAX).

## Data accessibility

All newly generated data have been deposited in GenBank (accession numbers pending) and the code is available on GitHub (https://github.com/ValentinaBoP/Wevolution).

## Author contributions

V.P. and A.S. designed the study. V.P. analysed the data and wrote the first manuscript draft. V.P. and A.S. wrote the subsequent drafts and all authors revised the subsequent drafts. M.I. and A.S. conducted laboratory work. C.F.-V., B.M., H.A., C.T., B.W.B., L.C., M.G., and M.I. collected and provided samples. M.P.K.B. and M.I. established the birds-of-paradise taxon sampling and provided unpublished genomic data.

